# Validation of genetic variants from NGS data using Deep Convolutional Neural Networks

**DOI:** 10.1101/2022.04.12.488021

**Authors:** Marc Vaisband, Maria Schubert, Franz Josef Gassner, Roland Geisberger, Richard Greil, Nadja Zaborsky, Jan Hasenauer

## Abstract

Accurate somatic variant calling from next-generation sequencing data is one most important tasks in personalised cancer therapy. The sophistication of the available technologies is ever-increasing, yet, manual candidate refinement is still a necessary step in state-of-the-art processing pipelines. This limits reproducibility and introduces a bottleneck with respect to scalability. We demonstrate that the validation of genetic variants can be improved using a machine learning approach resting on a Convolutional Neural Network, trained using existing human annotation. In contrast to existing approaches, we introduce a way in which contextual data from sequencing tracks can be included into the automated assessment. A rigorous evaluation shows that the resulting model is robust and performs on par with trained researchers following published standard operating procedure.

## 1 Introduction

Over the last decades, extensive research has been conducted to unravel the molecular, cellular and immunological mechanisms involved in cancer development. Especially the extensive use of next-generation sequencing (NGS) technologies enabled a progressively more cost-effective approach for the discovery of genetic variants within the genome that are inherited (germline mutations) or acquired (somatic mutations) and thereby predispose for or contribute to cancer development. So far, a huge amount of NGS data has been systematically analysed to determine meaningful information on potential drivers of tumourigenesis, dynamics of tumour evolution and differences between tumour entities regarding average mutational burden, tumour-specific mutational signatures or genetic diversification contributing to drug resistance and disease relapse [1, 2, 3, 4].

In personalised cancer therapy, a common task is to identify somatic mutations from paired samples of tumour and normal tissue of individual patients to select appropriate treatment options. In a typical processing pipeline for tumour-normal pairs, the sequenced reads for each sample are aligned to a reference genome, before undergoing removal of duplicate fragments and base quality score recalibration (BQSR). This process yields NGS data ready for further analysis (most commonly stored in the binary “BAM” file format), and the normal (“germline”) can be compared to the tumour sample to obtain somatic mutations in a process known as variant calling.

Due to the importance of variant analysis in modern biomedical research, the field has seen the emergence of a number of variant calling algorithms. These range in their methodologies from classical bioinformatical approaches (e.g. VarScan2 [47], Strelka2 [6] or SomaticSniper [7]) to deep learning, notably the seminal DeepVariant [29] and others like Clairvoyante [9] or NeuSomatic [10]. Despite comprehensive large-scale benchmarking, no single configuration of tools has emerged as the approach of choice for all settings, with them all having respective advantages and drawbacks (see e.g. [11, 12, 14]). It is currently common practice to cross-reference predictions made by several different variant callers [14] [15]. Regardless, any variant analysis workflow must currently still incorporate a manual refinement of candidate variants following published standard operating procedure by trained researchers [16, 59, 18] (for a broad overview of a typical variant calling workflow, see Figure 1). Besides being immensely time-consuming, this is intrinsically at odds with the principles of reproducible research, as there is by necessity a grey area where different researchers may come to different conclusions when examining a candidate variant, despite following the same guidelines (Barnell et al. [59] give a figure of 94.1% for the accuracy of reviewers following the proposed procedure). Coupled with the fact that different pipelines may yield very different results, this points to a great need for uniformisation, something which has been clear since the advent of NGS [19]. Very recent studies, too, point to the immense need for automation in the field, as the huge diversity in variants makes manual curation infeasible [20].

**Figure 1:**
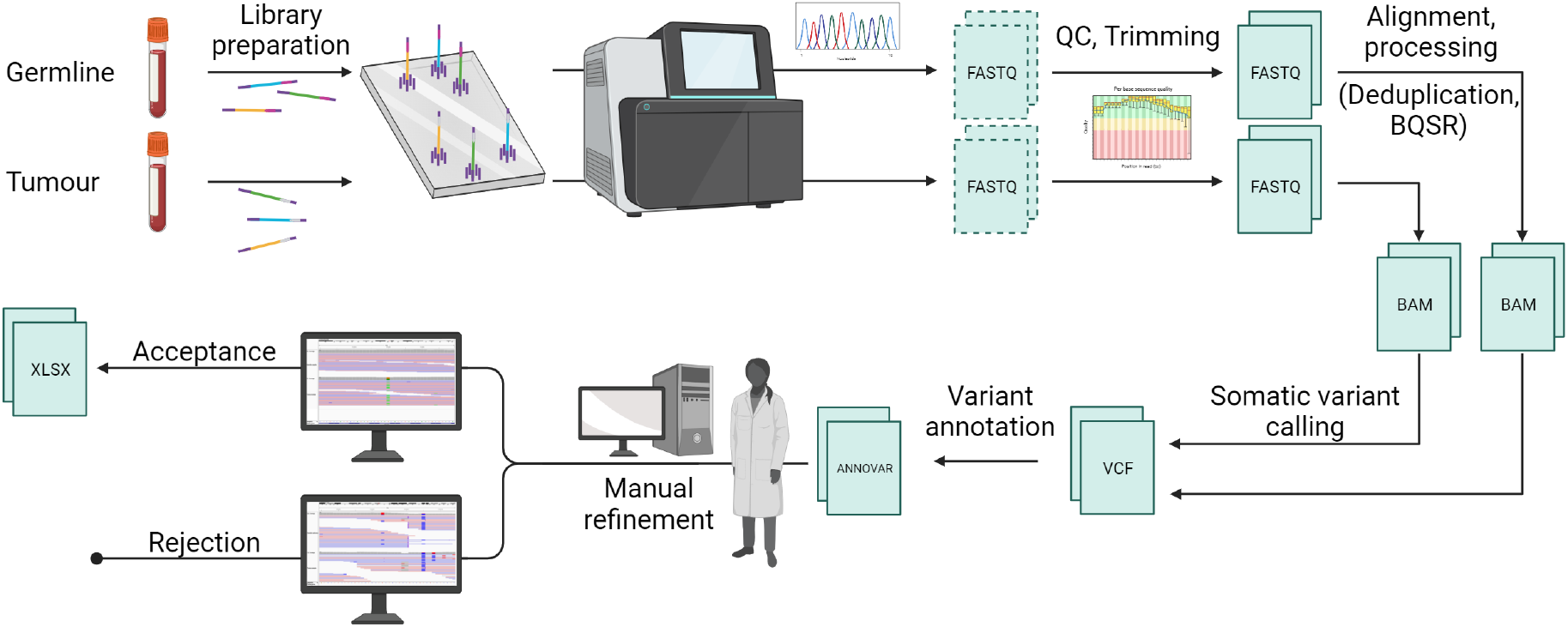
A typical processing pipeline from blood sample to somatic variant list.

One of the reasons for the need for manual refinement is that available pipelines still struggle with sequencing artefacts. The vast improvement in sequencing throughput capacity and cost-efficiency brought about by NGS has had a principal trade-off in reliability and accuracy (see e.g. [21, 22]). This bargain has been whole-heartedly embraced by the majority of the research community; it means, however, that sequencing artefacts are a common occurrence and must be accounted for when designing analysis pipelines. The reasons for this on a technological level are varied, and errors can be introduced at any step of the sequencing process. During library preparation, factors such as differential amplification, polymerase errors or the method and chemistry of DNA fragmentation can contribute to the generation of artefacts. Their recognition by a human reviewer rests on a number of factors such as low sequencing coverage, misaligned reads, strand bias, low base and alignment qualities and sequencing errors around the locus, especially present in regions of low complexity (Figure 2). Further artefacts may arise from sequencing during cluster amplification, cycle sequencing or image analysis [23, 24, 25, 26, 27].

**Figure 2:**
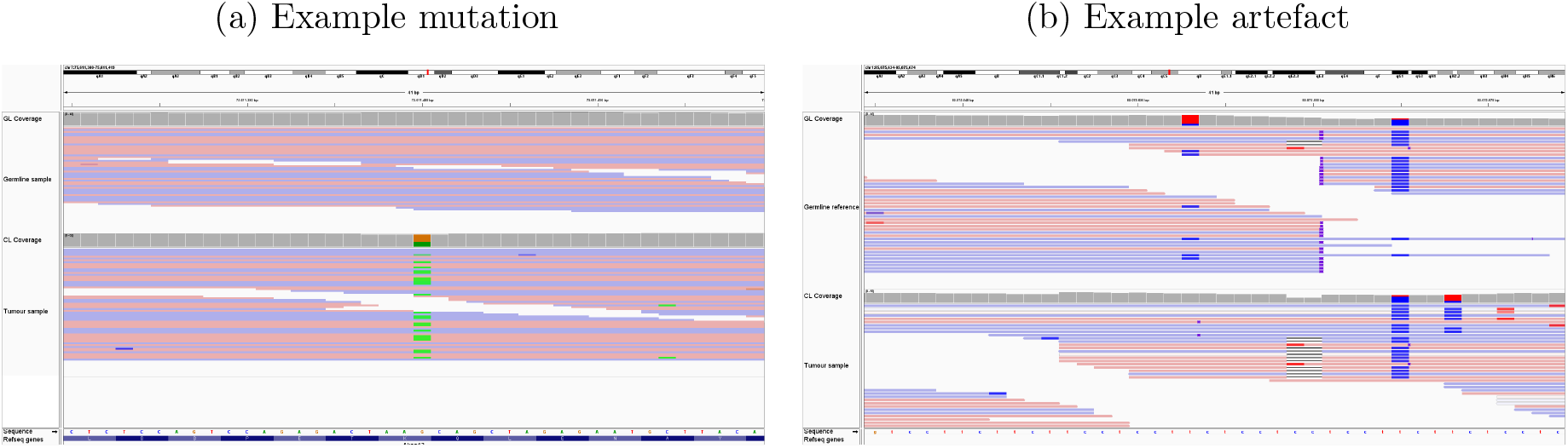
Examples for sequencing artefacts. Screen captures from the Integrative Genomics Viewer (IGV) [28]; in each case the germline track is shown in the upper half, and the tumour track in the lower. Shown are typical examples of (a), a genuine mutation, and (b), a sequencing artefact flagged as a candidate variant. The colour of the reads indicates their direction, unless they disagree with the reference genome, in which case the disagreeing base is coloured in. The artefact can be recognised by the low complexity of the DNA sequence and the subsequently misaligned abruptly ending reads, as well as general sequencing “noise”. entirely, germline and tumour samples would always have to be sequenced with the same library preparation. In longitudinal studies or transplantation experiments in particular, this would necessitate a new library preparation and full reanalysis of germline samples for each transplantation. In most applications, this is infeasible and prohibitively costly.

Artefacts can have many different sources, including library preparation. If the library preparation is done at the same time for both germline and tumour, this poses no problem, as the incorrectly called base will be present in both samples and no artefactual candidate variant is generated. If, however, the library preparations of germline and tumour sample differ, this can result in calls which appear highly credible at first glance. They can only be identified by consulting other, otherwise unrelated, samples which have been sequenced using the same library preparation. Coming across these artefacts is quotidian in research practice and manual variant refinement (see appendix 2 in the supplement for two examples), but was until now, to our knowledge, not implemented in any refinement tool. To exclude this error source Several tools have been published in the past to harness machine learning for variant calling and variant refinement. Yet, many such variant callers (including e.g. DeepVariant [29]) compare to reference genomes instead of employing tumour-normal pairs. This issue is addressed some variant refinement methods, but these methods do not exploit the benefits of deep learning. Ainscough et al. [30] present a case where a one-layer perceptron network was applied to manually defined summary statistics extracted from data, and a similar approach based on a Random Forest Classifier was also proposed by Li et al. [31]. The reliance on hand-crafted features extracted from sequencing data is a severe limitation compared to learned features, which can take better advantage of complex structure inherent to the data. Furthermore, none of the available methods exploit sequencing tracks besides the tumour-normal sample pairs, even though this can provide crucial information about sequencing artefacts in a specific experiment.

In this study, we propose a deep learning method for the automatic refinement of variant lists provided by variant calls. It employs convolutional neural networks, are a class of artificial neural networks that rely on “learning” (i.e. optimising with respect to a loss function) convolutional filters, which are applied across spatially arranged data. This makes them excel at tasks which are informed by local structure. They were brought into the spotlight of machine learning research by the immensely influential work of Krizhevsky, Sutskever and Hinton in [32] in the field of image classification. Since this renewal of interest, they have been used to great success in many fields, among them image processing, biomedical applications, and intersections between the two (see for example [33, 34, 35, 36, 37] and the citations contained therein). We demonstrate that the proposed method achieves a high degree of accuracy on different datasets.

## Methods

In this section, we introduce the deep learning method for variant refinement. Its inputs are BAM files obtained by sequence alignment, VCF files obtained by variant calling, and metainformation (e.g. on library preparation). The output is a list of refined variants from which sequencing artefacts have been removed. Below, we discuss the considered dataset and the evaluation of the method.

### 1.1 Input data representation

The application of deep learning methods for variant refinement requires a numerical representation of the sequencing data. In this study, we encode the input data as a three-dimensional tensor, with the dimensions corresponding to

- position of the base on the reference genome,
- index of the read, and
- base-wise information (nucleotide type, quality scores, and read direction).

We consider a fixed symmetrical window of d_window_ bases around the potential variants listed in the VCF files provided by variant callers and a maximum of d_reads_ reads. Base-wise information are the nucleotide type, base-wise quality, read alignment quality, as well as a binary flag showing whether a read was reversed. For the nucleotide type, a one-hot encoding is employed, resulting in four entries (one for each nucleotide), as is typically the case in Deep Learning with sequencing data [38, 39, 40]. Overall, we have seven entries per base – but this can be extended by further input features if desired. Whenever a read did not cover a position in the base window, or there were fewer than d_reads_ reads available, the corresponding positions in the tensor were zero-padded. Overall, this process yields tensors in 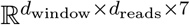.

To assess the presence of absence of a variant at a specific locus, we apply this encoding scheme to the sequencing data from the tumour and the normal tracks, as well as randomly chosen sequencing tracks with no biological relation to the variant (i.e. a different transplantation line), but the same library preparation. The additional tracks are used to provide context information, e.g. on library preparation, etc.

The tensors representing the information of the different sequencing tracks are concatenated along the third (i.e. “depth”) dimension. The resulting structure can be thought of as similar to the visualization researchers would see when inspecting a candidate with a genome viewer with different sequencing tracks aligned alongside each other, except that our data encoding forms a three-dimensional tensor where identical positions in the genome are aligned, cf. Figure 3.

**Figure 3:**
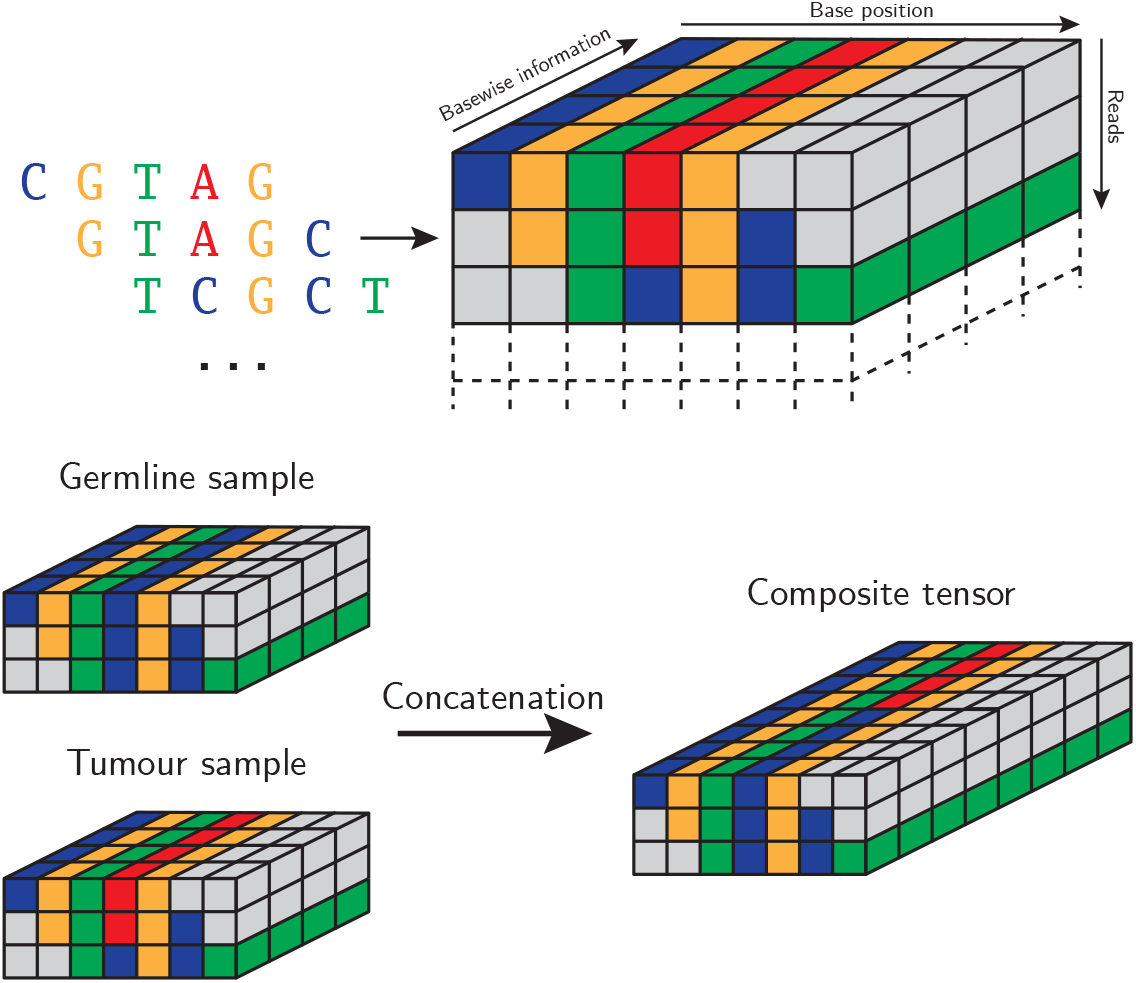
Process for obtaining composite data points from several sequencing tracks. Additional context tracks are added by the same principle.

### 1.2 Model topology and training

For the variant refinement, we employed a feedforward neural network incorporating convolutional layers. This type of topology has proven valuable for various sequence analysis tasks [41]. We evaluated different structures and observed a high degree of robustness of the results. For the subsequently presented results, the precise structure is as follows:

- Three convolutional blocks with filters of size 3 × 3 each (and ReLU activation). Using 32, 16 and 32 filters, respectively, each followed by Max-Pooling with the same filter size.
- One layer of 1 × 1 convolutions with 32 filters and ReLU activation
- A batch normalization layer with a momentum parameter of 0.8 followed by flattening
- A dense layer with 10 neurons and ReLU activation, and dropout of probability 0.2 applied on the out-edges for training
- A classification layer with two neurons and softmax activation

A visual summary is provided in Figure 4.

**Figure 4:**
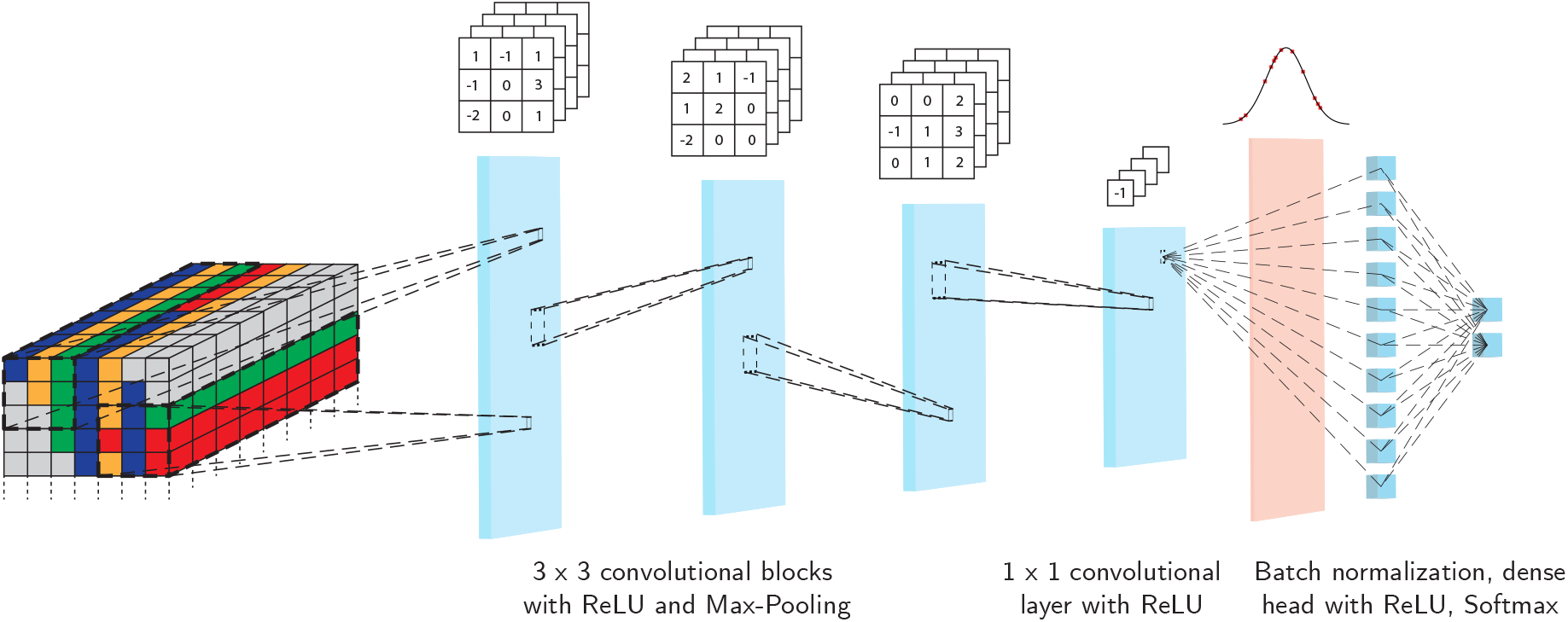
Architecture of neural network for classification. Input data runs through blocks of 3×3 convolutions with ReLU activation and max-pooling, before weighted averages over the filters are taken using 1 × 1 convolutions with ReLU. The results are flattened, normalised and passed to a dense classification head, where dropout is applied for regularisation.

Notable features of the selected architecture are the relatively small numbers of filters, and the lack of a large dense classification head which has been replaced by a 1 × 1 convolutional layer, “forcing” complexity into the filter structure. The overall number of trainable parameters, 27, 312, is low by neural network standards, which helps avoid excessive overfitting.

For training, we chose the binary cross-entropy loss function, as is most frequently done in binary classification. For further regularization, the training labels were subjected to *label smoothing* with smoothing parameter 0.1, meaning that the categorical (1, 0) and (0, 1) labels were converted into (0.95, 0.05) and (0.05, 0.95) respectively. Finally, training was done with the Adam optimiser [56].

### 1.3 Datasets

For the evaluation of the method we considered different datasets, focusing exclusively on candidates for single nucleotide variants (SNVs). The first dataset, generated in-house, contains 2085 candidate variants and is used for training and cross-validation. The second encompasses 1652 candidate variants and was obtained from data published in [61]. Only in-house data was used for training, with the entire out-of-facility dataset reserved for validation.

#### In-facility

To generate a first data set, we performed WES sequencing for murine samples of TCL1 and TCL1-AIDKO primary and transplanted tumours generated with the Agilent SureSelect XT Mouse All Exon Kit[48, 52]. This experimental system is a well-established widely for studying chronic lymphocytic leukemia (CLL) [46] and provided us with a highly controlled data set. The dataset includes 11 germline (GL) and 40 CLL samples (11 primary and 29 serially transplanted), based on which 40 comparisons were performed using VarScan2 [47], in order to identify somatic mutations occurring between germline and primary tumour, or transplanted tumour sample. WES data are accessible on Sequence Read Archive, NCBI, NIH (BioProject: PRJNA475208 [48], BioProject: PRJNA725403 [52] and BioProject: PRJNA789482. The variant candidates provided by VarScan2 were evaluated by three experienced researchers following published Standard Operating Procedure [59] to provide a ground-truth.

#### Out-of-facility

As a second data set, the results of a mutational analysis by Kotani et al. investigating WES data from murine progenitor cells transduced with human *MLL/AF9* and differentiated into bone marrow cells as a model for MLL-rearranged acute myeloid leukemia (AML) were used. Similar to the in-facility samples, primary and serially transplanted tumour samples were prepared and sequenced with the Agilent SureSelect XT Mouse All Exon V2 Kit [61]. The dataset encompasses 42 paired GL and AML samples (8 primary and 34 transplanted). Candidate variants were generated and manually annotated while cross-referencing with the list of mutations published with the data.

Details on the exact bioinformatics processing pipelines for both datasets are given in appendix 1 in the supplement.

### 1.4 Evaluation

We employed several strategies to assess the proposed method’s generalisation performance. On the in-house dataset, we used stratified 5-fold cross-validation, as well as repeated random stratified train-test splits. Additionally, the out-of-facility dataset was used as a hold-out validation set.

In all cases, the model’s aptitude was assessed using the standard metrics for binary classification tasks: accuracy, recall (also known as *sensitivity*), precision, and F1 score. If we let *TP*, *FP*, *TN*, *FN* denote true positives, false positives, true negatives, and false negatives, respectively, we define

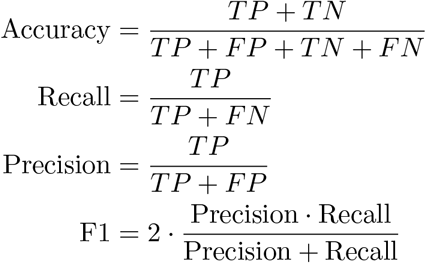

To gain further insights, we examined whether its accuracy on subcategories of the dataset differed significantly from the overall accuracy. In particular, we investigated possible differences in performance by reference base, alternative base, reference-alternative-pair, mutation class, and variant allele frequency (VAF). In each case, we used a binomial test to assess whether the null hypothesis that all results on subsets of the data have been obtained using the same accuracy can be rejected. The results were corrected for multiple testing using the Bonferroni method. For an overview of the overall data processing pipeline, see Figure 5.

**Figure 5:**
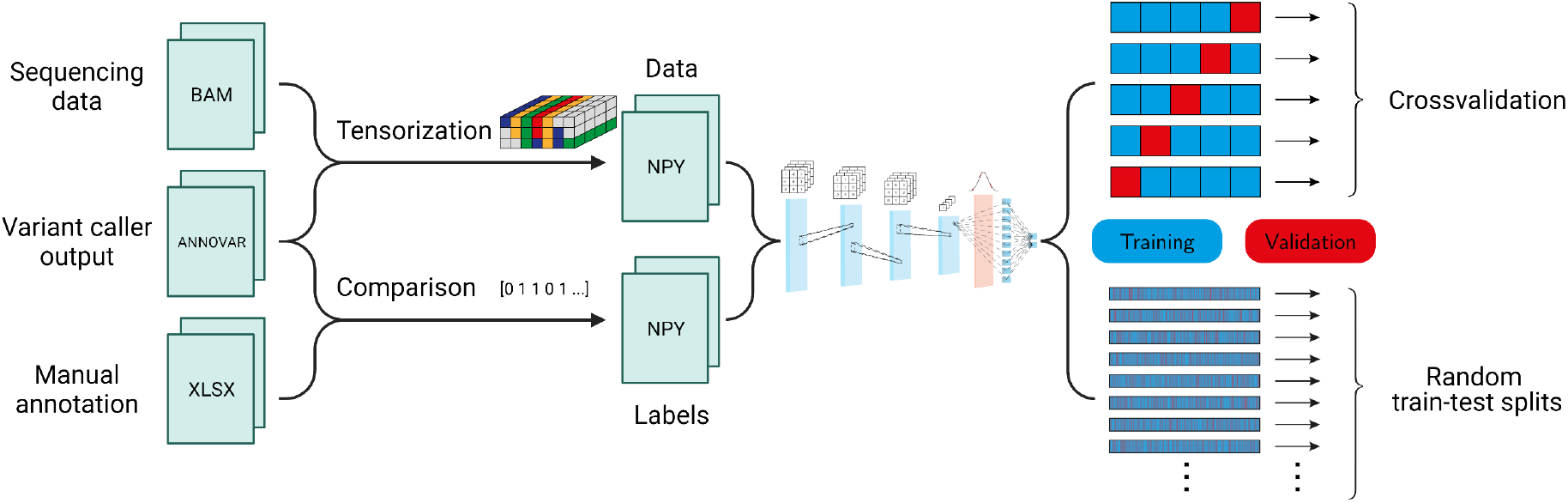
Overall model training and evaluation pipeline. The sequencing data is tensorised at candidate loci, while previous expert annotation provides labels. This dataset is then used to estimate the proposed neural network’s generalisation ability using two evaluation approaches.

## 2 Implementation

We performed the mutational analysis using the pipeline described by Schubert et al. [52]. Somatic variant calling using VarScan2 was performed using relatively permissive parameters, i.e. min-coverage-normal 5, min-coverage-tumour 5, min-var-freq 0.05, somatic-p-value 0.05 and strand-filter 1, and filtering of high confidence calls was performed according to Basic Protocol 2 published by Koboldt et al. [53], to reduce the false negative rate. The precise setting of VarScan2 dependent on the dataset, using internal standards of the clinical groups in Salzburg for the in-facility dataset and published parameters for the out-of-facility dataset (see Supplementary Table 2). The resulting variant lists were filtered for the following mutation classes (coming from ANNOVAR analysis “Func.RefGene”): “exonic”, “ncRNA_exonic”, ‘‘splicing”, “UTR3”, “UTR5”, “UTR5;UTR3”, ‘downstream”, ‘upstream”, “upstream;downstream”].

We implemented the proposed variant list refinement method using the Keras [54] interface to the TensorFlow [55] framework in Python. For training, we chose the canonical binary crossentropy loss function, and performed optimization with Adam [56] using a constant learning rate of 10^-3^. Optimization steps were done in batches of 256, for 50 epochs. The evaluation was done using the implementation of stratified cross-validation and train-test-splits provided by the scikit-learn [60] toolkit for Python. For the binomial test between data subsets, we chose the implementation in the statsmodels package for Python [58].

For the tensor representation of the data, we chose a window size of d_window_ = 101 bases around the examined locus. Reads were considered up to a maximum number of d_reads_ = 200 reads from each track. To handle potential homologies distorting the reported classification performance, we base-scrambled all input data, applying random A-T and/or C-G base swaps to each data point.

The complete implementation is available on GitHub (https://github.com/marc-vaisband/deepCNNvalid).

The final version used for the study is archived at Zenodo (https://zenodo.org/record/6409366).

## 3 Results

In the following, we illustrate and evaluate the accuracy of the proposed variant filtering approach. For this purpose, we consider the afore-described in- and out-of-facility datasets.

### 3.1 Information on sequencing context improves classification efficiency

To evaluate the proposed method and to understand the dependence of its performance on different factors, we first studied the in-house dataset. For this the precise experimental setup is well known and the use of mouse data from several transplantation rounds provides us with additional controls (e.g. an the plausibility of the absence and presence of mutations). We processed the data as described in the *Implementation* section. This yielded 2085 data points (of them 703 mutations and 1382 artefacts), each corresponding to a particular tumour-normal sample pair and a position in the genome. To generate a ground truth datasets, the list of variant candidates was screened by three experienced researchers following the guidelines by Barnell et al. [59]. The consensus annotation by these researchers provided the corresponding labels, dividing the candidates into true mutations and artefacts.

In a first step, we performed stratified 5-fold cross-validation, using the scikit-learn [60] implementation, with a fixed random seed, splitting the in-house dataset into five folds. We find that the proposed method, which accounts for context, achieves a good performance as measured in validation accuracy, validation precision, validation recall and validation F1 score, averaged over the five stratified folds. To assess whether the method can be simplified by removing context information, we reran the evaluation with reduced input tensors and neural network. This revealed that context information indeed improves the validation performance, by 0.02 points compared to the context-free case. Most likely, this is due to added information on the overall quality of a sequencing run and sequencing artefacts.

| Dataset | Accuracy | Precision | Recall | F1 |
| --- | --- | --- | --- | --- |
| With context tracks, average | 0.968 | 0.953 | 0.954 | 0.953 |
| Without context tracks, average | 0.952 | 0.936 | 0.92 | 0.927 |

As the training time of the model is moderate (408 seconds on a node with 56 CPUs and no GPU acceleration), we assessed the reliability of the results by considering 100 random stratified train-test-splits of our data. We again observe that including context tracks improves the performance metrics, by 0.1 to 0.2 points.

| Dataset | Accuracy | Precision | Recall | F1 |
| --- | --- | --- | --- | --- |
| With context tracks, average | 0.958 | 0.934 | 0.945 | 0.937 |
| Without context tracks, average | 0.946 | 0.921 | 0.923 | 0.920 |

A visual summary of both evaluation results can be found in Figure 7; the detailed crossvalidation scores per fold are included in the appendix 3 found in the supplement.

**Figure 6:**
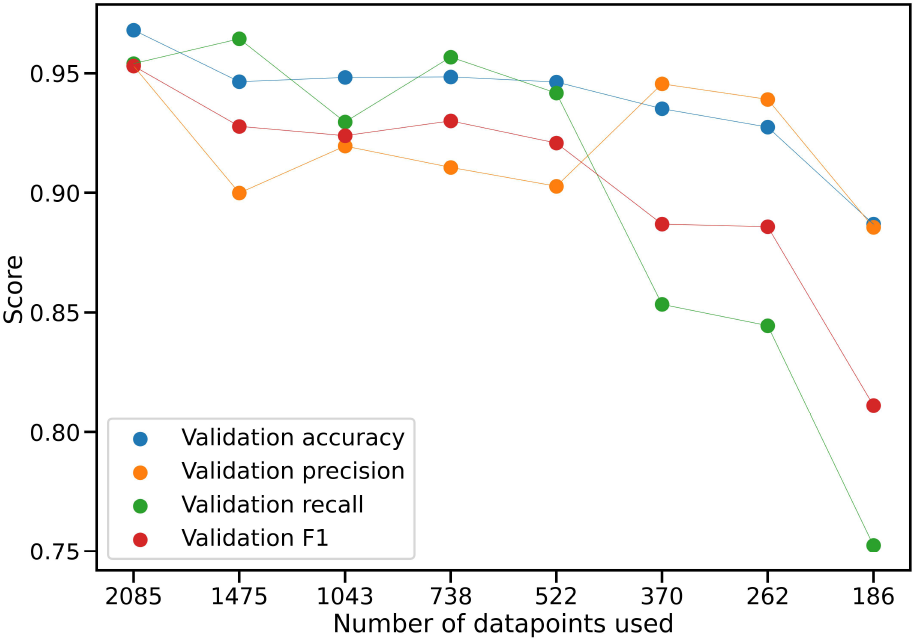
Results of data subsampling. The scores of successive cross-validation runs after repeatedly sub-sampling the data with a factor of 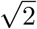.

**Figure 7:**
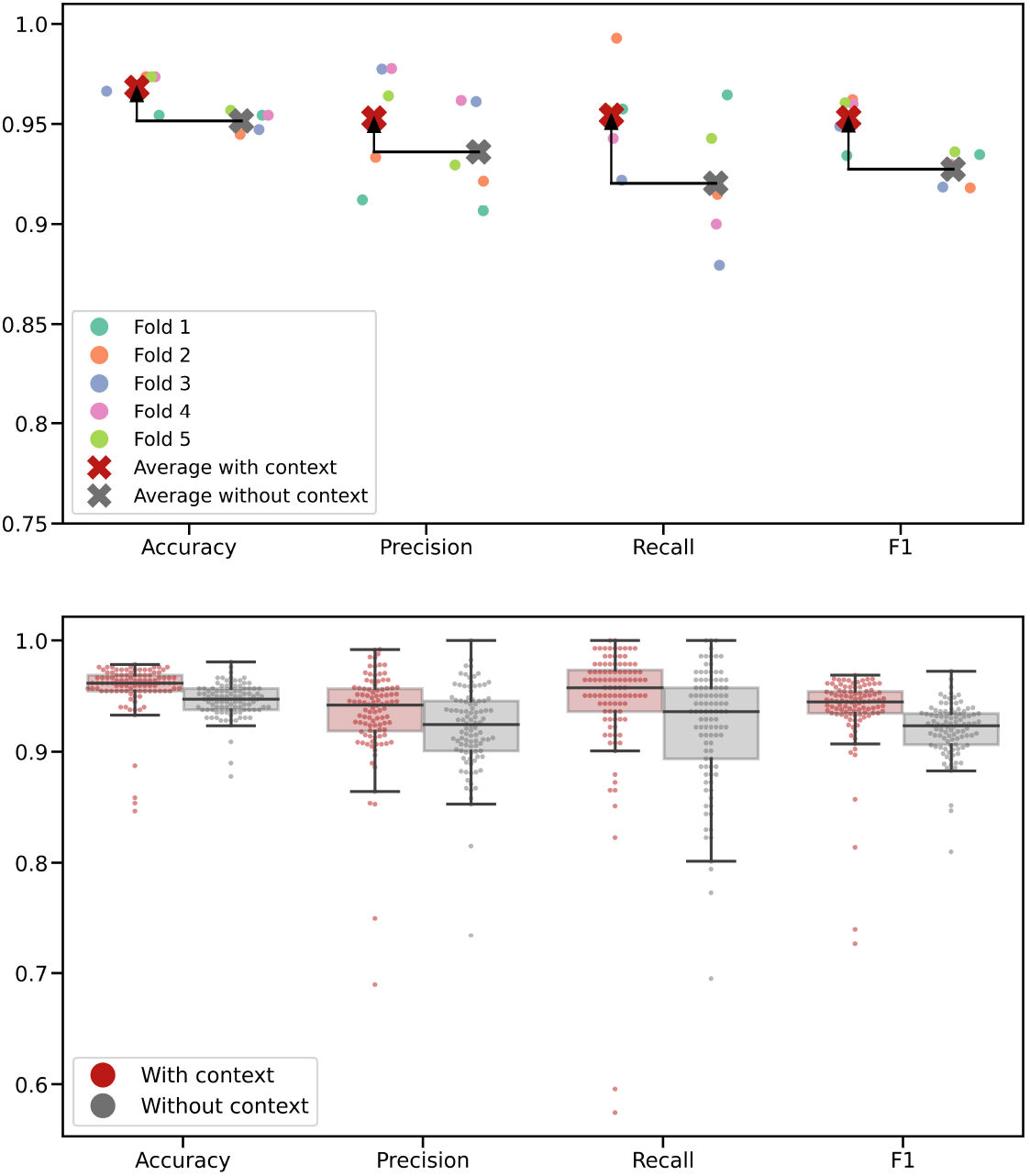
Results of model evaluation. Modelling with context tracks shown in red; without in grey. In the top figure, the results of 5-fold cross-validation are shown, with arrows to indicate the improvement achieved by including context. In the bottom figure, the results of 2 × 100 optimization runs with random train-test splits are presented.

### 3.2 Variant candidate refinement performs well even for limited training set sizes

As the proposed method performed rather well despite the relatively limited number of training examples provided by the in-house dataset, we assessed the dependence of the validation performance on sample availability. Therefore, we progressively sub-sampled the training set, repeatedly reducing its size by a factor of 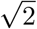, and performed 5-fold cross-validation on each subset. We observed that as expected, overall performance declines as the number of data-points becomes smaller, but that this decline is initially relatively gradual, before becoming more precipitous as the sample size dips below 500 (cf. Figure 6). This supports the conventional wisdom that deep learning methods often need sample sizes well into the hundreds in order to perform well.

### 3.3 Classification performance does not depend on bases or mutation class

For the potential use of the proposed method in clinical practice, it is crucial to gain a precise understanding of the model’s behaviour with respect to different kinds of candidate variants. In particular unbiasedness is an important feature. To analyse this, we subdivided the in-house data based on reference base, alternative base, and mutation category. Using the validation predictions from the cross-validation, we investigated whether the validation accuracy on any one group based on these distinctions differed significantly from the overall accuracy, using a binomial test. No significant differences were found. This suggests that the model performs equally well in all cases - irrespective of the reference and alternative bases, as well as mutation category.

### 3.4 The model generalises to independent out-of-facility data

As the model perform well on the in-facility data, we performed the arguably most rigorous, assessment of generalisation ability for any machine learning model is of course its application, and tested it on data which was generated completely independently of the data used for training or even in-facility validation. For this purpose, we utilised the data published in [61] and processed as outlined in the *Implementation* section.

We trained a model on all 2085 of our in-facility data points, and evaluated it on the 1652 candidate variants from the out-of-facility data.

The final evaluation of the out-of-facility dataset for independent validation yielded an accuracy of 0.949, a precision of 0.948, a recall of 0.995, and an F1 score of 0.971. These levels are similar to the in-facility data used for training, showing that the method generalizes to different sequencing methods and experts.

## 4 Discussion

In this manuscript, we considered the identification of somatic variants from NGS data, which is one of the key challenges in modern cancer research. It is aided by ever-improving sequencing and variant calling technology, but currently still dependent on manual candidate refinement, which presents a huge opportunity for improvement in terms of scalability and reproducibility. Here, we demonstrate that Convolutional Neural Networks can provide accurate classifiers and proposed a concrete network topology.

Our results demonstrate that the proposed method is indeed well-suited for the task. It features low validation error rates in cross-validation and generalises equally well to independent data from a completely different facility. The achieved performance is in both cases on par with specifically trained human researchers following standard operating procedure. Moreover, our analysis shows that the inclusion of additional sequencing tracks for added context improves classification results by a considerable margin - while the improvement in percentage points is small due to the high overall accuracies in both cases, the error rate is cut by roughly a third.

A similar approach would likely be equally successful for the refinement of germline variants, which have recently been the subject of particularly active research in cancer biology ([62], [63]). Here, too, the recognition of technical artefacts presents a formidable challenge [64]. The same methodology as presented in this work would allow a sequenced germline sample to be tensorised and evaluated by a neural network. In particular, it should be noted that the idea of including additional context tracks to handle library-specific artefacts translates analogously to this case, so that sequencing data of unrelated samples with the same library preparation would be added along the depth dimension.

Of course, further research is still needed in several directions. One natural avenue for exten sion would be the inclusion of indels, where repetitive sequences play a large role, so that we would once again expect a convolutional approach to do well. Another would be the inclusion of additional datasets for training and evaluation. It is unfortunately not common for lists of identified sequencing artefacts to be published, so building appropriate training data from public datasets is non-trivial. We hope, however, that this will change in the future, because any application of machine learning methods to sequencing data stands to profit greatly from the availability of this information. From a biological point of view, the most relevant followup would be a thorough investigation of the possible causes for library-specific sequencing artefacts; while they are routinely observed in practice, the authors are not aware of any comprehensive analysis of the underlying mechanisms.

To conclude, we have presented a method which can contribute to the standardisation of somatic variant calling and will benefit the research community by improving efficiency, reproducibility and interoperability. We hope that given time and data availability, tools such as this may become widely accepted in everyday research and clinical practice.

## Data availability

All code used in this publication is available at GitHub (https://github.com/marc-vaisband/deepCNNvalid), with the final version archived at Zen-odo (https://zenodo.org/record/6409366). The used WES data is available at SRA PRJNA475208, PRJNA725403 and PRJNA789482.

## Ethics declaration

Mouse experiments for the data used in this work were performed under the approval of the Austrian animal ethics committee (BMWF 66.012/0009-II/3b/2012, TGV/52/11-2012 and BMBWF-66.012/0002-V/3b/2018).

## Figure credit

Figures 1 and 5 were created using BioRender (https://biorender.com/).

## A Details on bioinformatics pipelines

**Table 1:** List of different versions of bioinformatics tools used for WES data analysis of in-facility training data set, out-of-facility validation data set. Tool versions used by Kotani et al. are not available. NA = not available; *v3.7 used for RealignerTargetCreator, IndelRealigner, DepthOfCoverage.

| Tools | In-facility | Out-of-facility | Kotani et al. (ref 21, 26) |
| --- | --- | --- | --- |
| BWA-MEM | v0.7.15 | v0.7.15 | NA |
| PicardTools | v2.2.2 | v2.21.9 | NA |
| GATK | v3.7 | v3.7*; v4.1.7.0 | NA |
| samtools | v1.3.1 | v1.10 | NA |
| VarScan2 | v2.4.2 | v2.4.4 | NA |
| ANNOVAR | v2015Dec14 | v2017July17 | NA |

**Table 2:** List of VarScan2 filter settings used for variant calling of In-facility training data set and Out-of-facility validation data set in house and list of settings used by Kotani et al. *p-value 0.05, if more than 20 percent of the mutated allele in tumour sample and less than 10 percent of the mutated in normal sample

| Variant calling | In-facility | Out-of-facility | Kotani et al. (ref 21, 26) |
| --- | --- | --- | --- |
| min-coverage-normal | 5 | 10 | 10/8 |
| min-coverage-tumor | 5 | 10 | 10/8 |
| min-var-freq | 0.05 | 0.07 | 0.07/0.05 |
| somatic p-value | 0.05 | 0.05 | 0.001 or *0.05/0.003 |
| strand filter | 1 | no | no |

## B Examples for library-specific artefacts

Below are two example positions which exhibit consistent sequencing artefacts across a library. Without this context, they could be mistaken for mutations.

**Figure 8:**
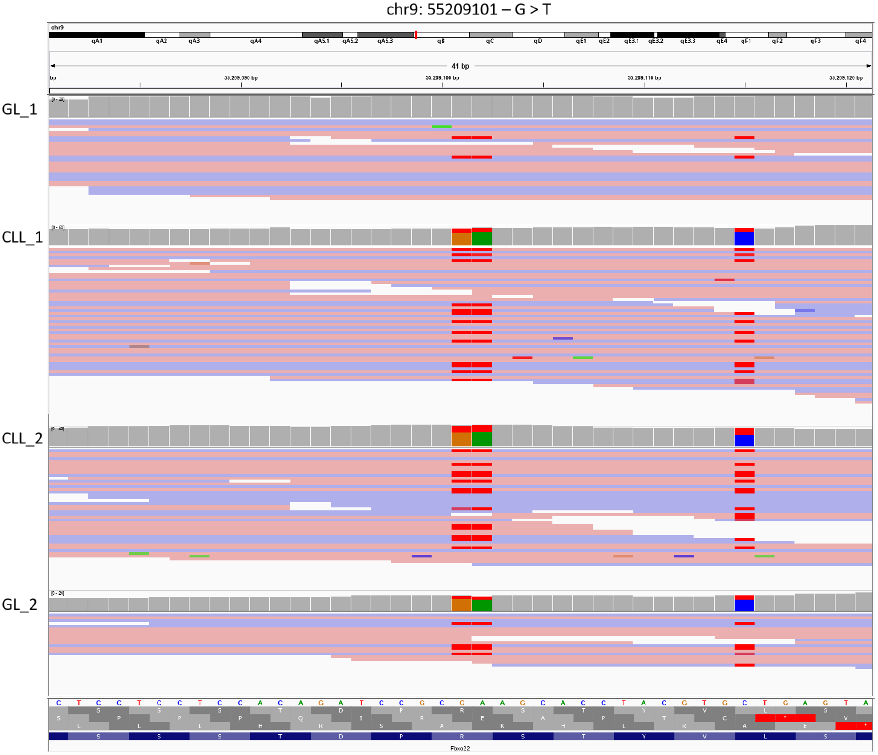
Example 1. SNV call from comparison of GL_1 (GL111) to CLL_1 (CLL111), but alteration recurs across library preparation, e.g. in a different tumour sample CLL_2 (CLL91), and also an unrelated germline sample GL_2 (GL91). Locus chr9, 55209101

**Figure 9:**
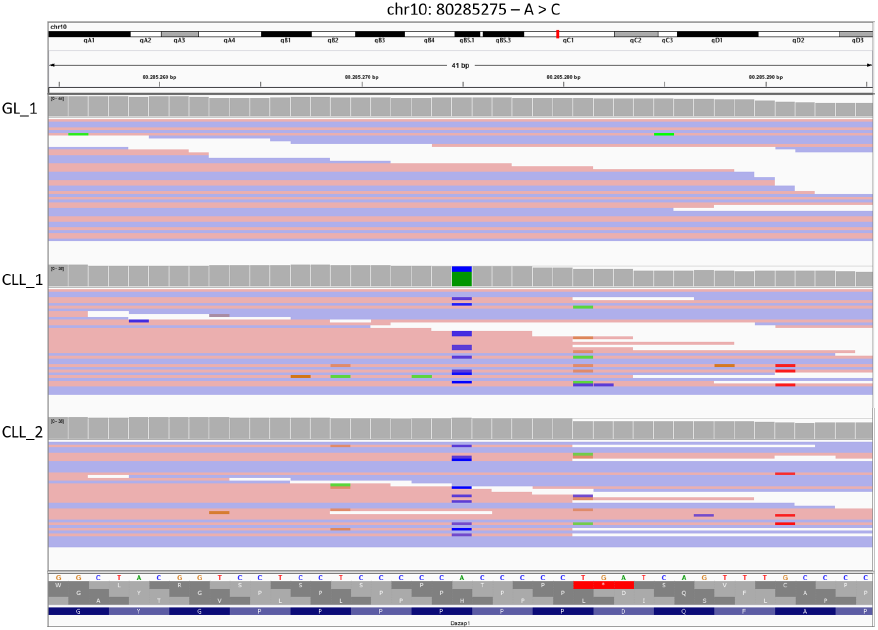
Example 2. SNV call from comparison of GL_1 (GL_C25) to CLL_1 (CLL6), but also recurs in the unrelated sample CLL 2 (CLL91). Locus chr10, 80285275

## C Cross-validation scores

Below, the individual fold scores for cross-validation with and without context tracks can be found. All values rounded to three decimals.

Without context:

**Table 3:** Detailed scores from 5-fold cross-validation with context tracks.

| Fold | Accuracy | Precision | Recall | F1 |
| --- | --- | --- | --- | --- |
| 1 | 0.954 | 0.912 | 0.957 | 0.934 |
| 2 | 0.974 | 0.933 | 0.993 | 0.962 |
| 3 | 0.966 | 0.977 | 0.922 | 0.949 |
| 4 | 0.974 | 0.978 | 0.943 | 0.96 |
| 5 | 0.974 | 0.964 | 0.957 | 0.961 |
| Average | 0.968 | 0.953 | 0.954 | 0.953 |
| Std. dev. | 0.008 | 0.029 | 0.026 | 0.012 |

**Table 4:** Detailed scores from 5-fold cross-validation without context tracks.

| Fold | Accuracy | Precision | Recall | F1 |
| --- | --- | --- | --- | --- |
| 1 | 0.954 | 0.907 | 0.965 | 0.935 |
| 2 | 0.945 | 0.921 | 0.915 | 0.918 |
| 3 | 0.947 | 0.961 | 0.879 | 0.919 |
| 4 | 0.954 | 0.962 | 0.9 | 0.93 |
| 5 | 0.957 | 0.93 | 0.943 | 0.936 |
| Average | 0.952 | 0.936 | 0.92 | 0.927 |
| Std. dev. | 0.005 | 0.025 | 0.034 | 0.009 |

